# QTL mapping and identification of corresponding genomic regions for black pod disease resistance to three *Phytophthora* species in *Theobroma cacao* L

**DOI:** 10.1101/366054

**Authors:** MA Barreto, JRBF Rosa, ISA Holanda, CB Cardoso-Silva, CIA Vildoso, D Ahnert, MM Souza, RX Corrêa, S Royaert, J Marelli, ESL Santos, EDMN Luz, AAF Garcia, AP Souza

## Abstract

The cacao tree (*Theobroma cacao* L.) is a species of great importance because cacao beans are the raw material used in the production of chocolate. However, the economic success of cacao is largely limited by important diseases such as black pod, which is responsible for losses of up to 30-40% of the global cacao harvest. The discovery of resistance genes could extensively reduce these losses. Therefore, the aims of this study were to construct an integrated multipoint genetic map, align polymorphisms against the available cacao genome, and identify quantitative trait loci (QTLs) associated with resistance to black pod disease in cacao. The genetic map had a total length of 956.41 cM and included 186 simple sequence repeat (SSR) markers distributed among 10 linkage groups. The physical “*in silico*” map covered more than 200 Mb of the cacao genome. Based on the mixed model predicted means of *Phytophthora* evaluation, a total of 6 QTLs were detected for *Phytophthora palmivora* (1 QTL), *Phytophthora citrophthora* (1 QTL), and *Phytophthora capsici* (4 QTLs). Approximately 1.77% to 3.29% of the phenotypic variation could be explained by the mapped QTLs. Several SSR marker-flanking regions containing mapped QTLs were located in proximity to disease regions. The greatest number of resistance genes was detected in linkage group 6, which provides strong evidence for a QTL. This joint analysis involving multipoint and mixed-model approaches may provide a potentially promising technique for detecting genes resistant to black pod and could be very useful for future studies in cacao breeding.

## Introduction

*Theobroma cacao* L. (also known as cacao or chocolate tree) is a perennial tree of the understory that belongs to the family Malvaceae. This tree is endemic to South American rainforests, which constitute a central region for the genetic diversity of these crops (Schultes and Cuatrecasas, 1964). *T*. *cacao* L. is a diploid species (2n = 2x = 20) (Davie, 1935) with an estimated genome length of approximately 430 Mb (Argout *et al*., 2011; Motamayor *et al*., 2013). Cacao beans are the raw material used in the manufacturing of chocolate and cacao butter, because this, crops have become an economically important commodity for more than 50 tropical countries, generating approximately 12 billion dollars in revenue yearly (ICCO, 2014).

Black pod (also known as *Phytophthora* pod rot, PPR) causes serious economic problems in all cacao-growing regions worldwide (Lanaud *et al*., 2009). This disease has affected cacao plantations since the 1920s and causes losses of up to 30-40% of the global production (Campêlo *et al*., 1982; Tahi *et al*., 2006). Black pod is caused by a complex of species belonging to the *Phytophthora* genus, which is known as the “plant destroyer” (Campêlo *et al*., 1982). Currently, seven *Phytophthora* spp. have been reported in cacao disease etiology: *P. katsurae* Ko and Chang, *P. megakarya* Brasier and Griffin, *P. megasperma* Dreschler, *P. citrophthora* RE Smith and EH Smith, *P. heveae* Thompson, *P. capsici* Leonian, and *P. palmivora* (Butler) Butler (Luz *et al*., 2004). Among these species, *P. palmivora* and *P. capsici* have a pantropical distribution, and both cause an estimated 20% to 30% annual loss and up to 10% of the tree deaths reported worldwide (Guest, 2007).

The symptoms and disease progression of black pod depend on the cacao genotype and the *Phytophthora* spp. involved; furthermore, they are influenced by climatic factors such as temperature, humidity and rainfall (Oliveira and Luz, 2005). Cacao pods are susceptible to fungal infection at all stages of development, and in the advanced stages of *Phytophthora* spp. infection, the beans become unsuitable for industrial use. Several measures have been used to control black pod disease, including appropriate cultural practices, fungicide application and the use of biological agents (*Trichoderma* spp.). However, these practices have substantial disadvantages, including increases in production costs, environmental pollution and ineffectiveness for field control (Nyassé *et al*., 2003). Thus, genetic resistance is of great importance as a more effective, economical and sustainable alternative for black pod control, and molecular markers have emerged as important tools in the search for more effective solutions for genetic control (Michelmore, 2003).

QTL mapping has been proposed for different crop species and for many complex traits including disease resistance (Kover & Caicedo, 2001; Clair, 2010) using almost all of the current classes of molecular markers. In general, numerous disease-resistance QTLs have been detected in plants, as reviewed in detail by Kover & Caicedo (2001). These mapped QTLs were important for the investigation of genomic regions that potentially contain candidate genes for disease resistance. QTL mapping has been specifically developed for the cacao tree for the identification of QTLs associated with resistance to *Phytophthora* spp. (Crouzillat *et al*., 2000a, b; Flament *et al*., 2001; Risterucci *et al*., 2003; Clement *et al*., 2003a; Lanaud *et al*., 2004; Brown *et al*., 2007).

However, none of the maps or linkage analyses published to date for cacao populations have been conducted using Wu’s multipoint approach (Wu *et al*., 2002; Tong *et al*., 2010). Unlike traditional two-point approaches, this procedure uses hidden Markov models (HMM) to estimate maximum likelihood based on the segregation patterns of all of the available markers in each linkage group, increasing the power to find the best ordering between them. The multipoint approach provides higher genetic information and increased saturation for the estimation of recombination fractions and linkage phases in map construction, which are conducted simultaneously in a full-sib population (Wu *et al*., 2002). Consequently, the search for QTLs according to their positions and genetic effects is also achieved in a multipoint context (Gazaffi *et al*., 2014), thereby increasing the power and confidence of the inferences.

Therefore, we propose that using a multipoint approach to construct the genetic linkage map will provide results with great power and confidence and enables conducting the QTL analyses for three *Phytophthora* species. The discovery of genomic regions containing resistance genes to black pod will be crucial for the inferring resistant phenotypes in future cycles of selection and for reducing the productivity losses resulting from this disease.

## Materials and Methods

No specific permits were required for the described field studies. This work was a collaborative project developed by researchers from the MCCS (Brazil and USA), USP (Brazil), UNICAMP (Brazil), UESC (Brazil) and UFOPA (Brazil), UESB (Brazil), and CEPLAC (Brazil).

### Plant Materials

The biological material used in the present study was obtained from the leaves of 265 individuals of an F1 population (full-sib family) derived from a cross between the heterozygous clones TSH 1188 (Trinidad selected hybrids; female parent) and CCN 51 (Coleccion Castro Naranjal; male parent). This population has been maintained in the MCCS located in Barro Preto, Bahia State, Brazil (14°42’45.021171” S and 39°22’13.008369” W). To obtain the F1 population, TSH 1188 and CCN 51 clones were maintained under controlled pollination according to the following procedure: the female flowers were protected for 24 h before pollination to avoid pollen contamination, and the cross was performed manually. The pods were collected and identified, the seeds were germinated, and the seedlings were planted in 3 × 3 m plots containing rows of 25 plants.

Both clones were selected because of their important contrasting agronomic traits, which include productivity, sexual incompatibility and disease resistance. Moreover, these clones are included in an international research program that is comprised of institutions from Brazil, Costa Rica and the United States. TSH 1188 was developed in Trinidad from the crosses of POUND 18 X TSH753 [open pollination X TSA 641 (SCA6 X IMC 67)] (ICGD, 2015), which produces rough red fruits, has self-incompatibility and is moderately resistant to black pod disease (Lopes *et al*., 2011). CCN 51 was developed by H. Castro in the early 1960s in Ecuador from the following crosses: (ICS 95 X IMC 67) X Oriente 1 (Boza *et al*., 2014); produces purplish-red immature fruits that become yellow-orange when ripe and have a slightly wrinkled rind, and the insides of the seeds have a clear purple color. This clone has been used as a parental genotype in many breeding and selection programs worldwide (Boza *et al*., 2014).

### Phenotypic Traits: Evaluation of Black Pod Disease

Phenotypic evaluation of black pod disease in the F1 population was performed separately for the species *P. palmivora*, *P. citrophthora* and *P. capsici*, as described previously by Barreto *et al*. (2015). These three species were used because they are predominantly found in cacao production areas in Brazil. The *Phytophthora* isolates were obtained from laboratory culture collections (Phytolab) belonging to CEPLAC (Comissão Executiva do Plano da Lavoura Cacaueira). Zoospore suspensions for each *Phytophthora* species were obtained from cultures grown on Petri dishes containing carrot-water (*P. citrophthora*) or carrot-agar (*P. palmivora* and *P. capsici*) media for at least 7 days (Suplementary Fig. 1).

Two 30-day inoculation series (trials) were conducted during the wet season for each *Phytophthora* species. In each inoculation series, 2-month-old leaves were collected early in the morning. Twenty discs from each individual were placed upside down on 8 plastic trays on 1-cm-thick dampened foam and incubated. Boxes containing a maximum of 48 cacao genotypes from the F1 population, parental clones TSH 1188 and CCN 51, and cultivars SCA 6 (resistant) and Catongo (susceptible) (Barreto *et al*., 2015) were assembled into five different sets of individuals (boxes) that were replicated four times. Each individual was represented by five discs within each box. The boxes were distributed throughout a small laboratory area under controlled conditions, and the leaf discs were not exposed to any light source. Symptoms were observed 7 days after inoculation using the 6-point scale of infection (lesions) proposed by Nyassé *et al*. (1995), where 0 = no symptoms; 1 = very small localized penetration points; 2 = small penetration spots, sometimes in a network; 3 = coalescing lesions of intermediate size; 4 = large coalescing brown patches; and 5 = uniform large dark brown lesions.

Since the study by Nyassé *et al*. (1995) was published, the leaf-disc test applied in this study has been widely used to screen for resistance to black pod disease in cacao trees in studies conducted by Barreto *et al*. (2015) and Bahia *et al*. (2015). This analysis provides a rapid and early assessment of resistance levels, furthermore, a positive correlation between the data obtained by this method and the data obtained by field analyses has been observed, as well as anatomical similarities between the abaxial leaf side and the cacao pod epidermis (Nyassé *et al*., 1995; Tahi *et al*., 2006; Santos *et al*., 2009). In addition, Magalhães, *et al*. (2016) used this method to realize an indirect screening approach for Ceratocystis wilt resistance and found a positive correlation between the leaf disc method and field resistance.

The phenotypic data obtained for each *Phytophthora* species, based on the 6-point scale of infection (lesions), were analyzed according to the following statistical model:

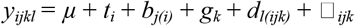

where *y_ijkl_* corresponds to the level of black pod infection; *µ* is the general average; *t_i_* is the fixed effect of the inoculation series (trial) *i*; *b_j(i)_* is the fixed effect of box *j* nested within trial *i*; *g_k_* is the random effect associated with genotype *k*; *dl(ijk)* is the random effect associated with disc *l* nested within genotype *k*, box *j* and trial *i*; and _*ijk*_ is the random residual term among plots. The analyses were performed using GenStat software v.13 (Payne *et al*., 2010) with a mixed-model approach (Henderson *et al*., 1959; Robinson, 2012).

Different structures of variances and covariances were investigated for the genetic and residual matrices of the described mixed model. Briefly, we assessed models that could account for the heterogeneity of variances or the presence of covariances (correlations) between observations. We used the restricted maximum likelihood (REML) method (Patterson and Thompson, 1971) to estimate the random components of the models. The Akaike information criterion (AIC) (Akaike, 1974) was used to compare and select the best model. The fixed effects were analyzed using the Wald (test) statistics. The predicted means for the individuals were extracted from the most likely mixed model and used for QTL mapping.

In addition, estimates of genotypic and phenotypic variances as well as estimates of heritability and coefficients of variation were obtained from the analysis of each *Phytophthora* species. Moreover, genetic correlations between each pair of *Phytophthora* species were estimated from the predicted means using the Pearson coefficient, and a global level of 5% was considered statistically significant. The correlation analyses were performed using R software (R Development Core Team, 2014).

### DNA Extraction and Polymerase Chain Reaction (PCR) Amplification

A modified *cetyltrimethylammonium bromide* (CTAB) method (Rehem *et al*., 2010) was used to extract total genomic DNA from the leaves of each individual of the F1 population and from the parental clones TSH 1188 and CCN 51. The DNA quality was evaluated on a 1% agarose gel and was compared with a standard lambda phage marker. The DNA quantity was estimated using a NanoDrop 8000 UV-VIS Spectrophotometer (Thermo Scientific, Brazil) at 260 nm.

Different SSR marker were available to genotype the F1 population, and the origin and institutions of these markers are described in Supplementary Table 1 (additionally, Appendix A and Appendix B present a detailed description of the loci). The PCR amplifications were performed in a Bio-Rad C1000TM Thermal Cycler (Bio-Rad, USA) with a 15-µL final volume containing 15 ng template DNA, 0.2 µM each primer (forward and reverse), 100 µM each deoxynucleotide (dNTP), 2.0 mM MgCl2, 10 mM Tris-HCl, 50 mM KCl, 0.25 µg µL-1 bovine serum albumin (BSA) and 0.5 units of Taq DNA Polymerase (Invitrogen, SP, Brazil). The PCR program included an initial denaturation at 95°C for 5 min, followed by 30 cycles at the appropriate melting temperature (Tm) for each primer for 1 min, 72°C for 1 min and 95°C for 1 min, with a final elongation step at 72°C for 30 min; the PCR amplification quality was evaluated on 3% agarose gels. Certain loci were subjected to electrophoresis on a 6% denaturing polyacrylamide gel in 1X Tris/Borate/EDTA (TBE) buffer, and a 10 bp ladder was used (Invitrogen, SP, Brazil) as a size standard. The DNA fragments were visualized using 0.2% silver-staining solution (Creste *et al*., 2001). Other loci were subjected to capillary electrophoresis in an ABI PRISM^®^ 3100 Genetic Analyzer (Applied Biosystems), an automated system used for the analysis of fluorescently labeled DNA fragments. GeneMarker^®^ software (SoftGenetics) was used to establish the peaks of filtering and interpretation, define the genotype of each individual, and generate the final compilation of data.

For each SSR marker, a classification (of 18 possible groups) was assigned to indicate the cross type and segregation (1:1:1:1, 1:2:1, 3:1, and 1:1) based on both parental and offspring marker band patterns, according to Table 1 proposed by Wu et al. (2002a).

**Table 1.** Characterization of the linkage groups and chromosomes from the multipoint linkage and physical maps.

| LG/Chr | Multipoint genetic linkage map (Figure 1) |  |  | Physical “*in silico*” map (Figure 2) |  |  | Proportion of the B97-61/B2 genome (%) |
| --- | --- | --- | --- | --- | --- | --- | --- |
|  | Length (cM) | Number of markers | Average distance between markers (cM) | Length (Mb) | Number of markers | Average distance between markers (Mb) |  |
| 1 | 135.75 | 30 | 4.53 | 30.02 | 30 | 1.00 | 77.00 |
| 2 | 115.35 | 23 | 5.02 | 26.64 | 30 | 0.89 | 62.77 |
| 3 | 91.76 | 22 | 4.17 | 24.44 | 30 | 0.82 | 71.05 |
| 4 | 108.38 | 27 | 4.02 | 22.69 | 26 | 0.87 | 67.75 |
| 5 | 124.77 | 22 | 5.67 | 25.31 | 27 | 0.94 | 62.59 |
| 6 | 74.67 | 11 | 6.79 | 14.90 | 14 | 1.06 | 54.58 |
| 7 | 61.87 | 13 | 4.76 | 13.61 | 15 | 0.91 | 55.83 |
| 8 | 71.72 | 07 | 10.25 | 9.07 | 07 | 1.30 | 42.12 |
| 9 | 114.86 | 24 | 4.79 | 27.69 | 30 | 0.92 | 65.87 |
| 10 | 57.28 | 07 | 8.18 | 8.37 | 08 | 1.05 | 32.88 |
| **Total** | **956.41** | **18.60** | **5.82** | **202.74** | **21.70** | **0.98** | **59.25** |

### Genetic Linkage Map and Genome Alignment

Marker segregation was assessed using a Chi-square test followed by the Bonferroni correction for multiple tests, according to the overall level of significance (α = 5%). The integrated genetic linkage map was constructed using OneMap software v.2.0-3 (Margarido *et al*., 2007) according to a multipoint approach based on the HMM (Wu et al., 2002a; Wu et al., 2002b). Initially, we obtained the pairwise linkage phases and recombination fractions between all the markers and separated these data into linkage groups (LGs) using a logarithm of the odds (LOD) score of 4.93 (an initial *empirical* threshold assigned based on the number of markers and Bonferroni correction) and a maximum recombination fraction of 0.30. Subsequently, the order of the markers within each LG was determined based on the HMM from a set of 5 initial markers, with the remaining markers subsequently added individually using the initial estimated order. The final multipoint recombination fractions between the markers were converted to centiMorgan (cM) units using the Kosambi mapping function (Kosambi, 1943). The final design of the map was generated using MapChart software v.2.2 (Voorrips, 2002).

The physical “in silico” map was constructed, which aligned the SSR marker sequences deposited in GenBank against the available cacao genome Criollo, using the nucleotide basic local alignment search tool (BLASTn) program. The cacao genome is available in the database *CocoaGen DB* (http://cocoagendb.cirad.fr), which has been developed and maintained by the Centre de Coopération Internationale en Recherche Agronomique pour le Développement (CIRAD) in France. An expected value (E-value) of 10 (e^-10^) was used to obtain an alignment with a lower probability of detecting false positives. An “*in silico*” PCR primer was used on the cacao genome v1.0 with an expected amplicon size between 40 to 1,000 bp, and a single mismatch was considered acceptable. The final design of the physical “*in silico*” map was also generated using MapChart software.

### QTL Mapping

QTL analyses were performed using the multipoint genetic linkage map according to the model proposed by Gazaffi *et al*. (2014). This approach is an extension of the composite interval mapping (CIM) of Zeng (1993) for an outbred population. Initially, the entire genome was scanned to detect QTLs, and the conditional probabilities of the QTL genotypes were calculated every 1 cM based on a specific interval between two flanking markers. A maximum of 20 cofactors were included in the model to control for QTLs outside of the intervals. The markers used as cofactors were included based on the stepwise procedure and AIC (Akaike, 1974) for the selection of the final model. A window size of 15 cM was used to control the underlying information from both sides of each interval. To determine the presence of QTLs, LOD score-based thresholds were determined using 1,000 permutation tests (significance level of 0.95) based on the method of Chen and Storey (2006). The proportion of phenotypic variation (*R^2^*) explained by each QTL was calculated using least squares estimation.

Subsequently, genomic regions containing evidence of QTLs were fully investigated and tested for three possible effects according to Gazaffi *et al*. (2014): (i) additive for one parent, (ii) additive for the other parent, and (iii) dominance (intra-locus interactions between the additive effects of both parents). Based on the significance and signal of the QTL effects, the linkage phase between QTLs and flanking markers and QTL segregation at 1:1:1:1, 1:2:1, 3:1 or 1:1 were inferred. Gazaffi’s CIM extension was performed separately for each of the three *Phytophthora* species.

## Results

### Genetic Linkage Map

To evaluate polymorphisms in the F1 population, 83 SSR markers with polymorphic patterns in the parental clones TSH 1188 and CCN 51 were amplified. Fifty SSRs (60.24%) exhibited good amplification results. Of these, 30 SSRs (60.00%) were polymorphic in the F1 population, which allowed for the identification of 84 different alleles. Other markers were also obtained from a database containing 199 SSRs from a group of various institutions involved in a cacao-breeding project – CIRAD, Universidade Estadual de Santa Cruz (UESC), Universidade Federal Rural do Semi-Árido (UFERSA) and MCCS. In total, 229 polymorphic SSRs were available for the construction of a genetic linkage map and for analyses of QTLs in the F1 population of the present study (Appendix A: mapped markers; Appendix B: unmapped/unlinked markers).

Of the 229 genotyped markers, 34 markers (14.85%) were heterozygous for TSH 1188, 48 markers (20.96%) were heterozygous for CCN 51, and 147 markers (64.19%) were heterozygous for both parents, exhibiting 2 (34.70%), 3 (54.40%) or 4 (10.90%) alleles. Two hundred twenty-nine Chi-square tests, followed by Bonferroni correction, were performed for the polymorphic loci. The results revealed that 89 (out of 95 – 41.48%), 48 (out of 52 – 22.71%) and 79 (out of 82 – 35.81%) loci exhibited an expected segregation of 1:1:1:1, 1:2:1 and 1:1, respectively. Thus, of the 229 markers, only 6 (2.62%), 4 (1.75%) and 3 (1.31%) markers showed significant deviations from the expected segregation of the respective classes, for a total of 13 (5.68%) deviated markers. However, because these markers did not show strong deviations of segregation, we used all of the information for the construction of a genetic linkage map. Three deviated markers (out of 13 – 23.08%) were incorporated into the final linkage map, and 10 (76.92%) remaining deviated markers were among the 42 (18.34%) SSRs that were not allocated into the map. Most of these unlinked markers presented a segregation of 1:1 (19; 45.24%), although a great proportion of segregation was also represented by 1:1:1:1 (16; 38.10%).

The multipoint genetic linkage map containing 186 SSRs is shown in Figure 1. Ten LGs were obtained from pairwise recombination fractions between the markers, which were considered linked when the estimated fractions were equal to or lower than 0.30 and when their LOD scores were equal to or higher than 4.93. We believe that the multipoint approach based on HMM provided accurate positioning of the markers and reliable distances from multipoint (upgraded) recombination fractions based on all of the available molecular information across each LG.

**Figure 1.**
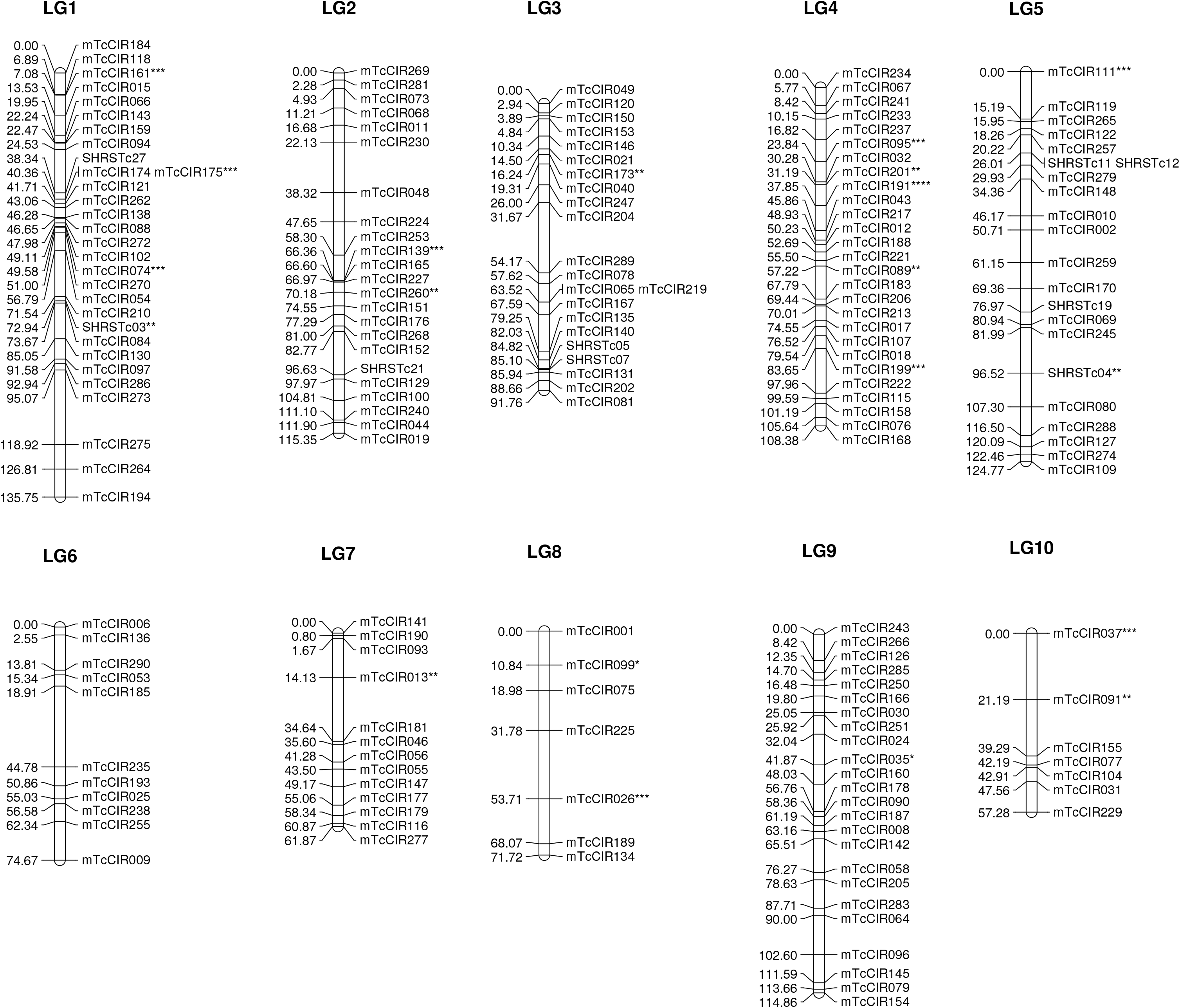
Multipoint integrated genetic linkage map constructed for the cacao tree (*Theobroma cacao* L.) using 265 individuals of an F1 segregating population (TSH 1188 X CCN 51). This map consists of 186 SSRs covering a total of 956.41 cM. The asterisks shown at the end of the name of some markers represent the following: *, deviation of segregation (ds); **, not aligned in the physical “*in silico*” map; ***, not aligned in the same group/chromosome of the physical map; ****, (ds) and not aligned in the physical map; and *****, (ds) and not aligned in the same group/chromosome of the physical map.

The total length of the multipoint linkage map is 956.41 cM (Table 1). Of the 10 identified LGs, corresponding to the haploid number of the cacao genome, six LGs exhibited larger genome coverage (in increasing order – LG 3, LG 4, LG 9, LG 2, LG 5 and LG 1). These LGs also had a higher number of markers (LG 3/LG 5, LG 2, LG 9, LG 4 and LG 1), varying from 22 to 30, and generally had lower average distances between two adjacent markers (LG 4, LG 3, LG 1, LG 9, LG 2 and LG 5), exhibiting a high level of saturation of the cacao genome.

### Genome Alignment: Physical “In Silico” Map

Recently, Argout *et al*. (2011) sequenced the genome of a suitable Belizean Criollo genotype (B97-61/B2), and these sequencing data are available in *CocoaGen DB*. The sequences of the 229 SSRs used in the present study were aligned against the B97-61/B2 genome. A physical “*in silico*” map was constructed from this alignment, and 217 SSRs were positioned to cover more than 200 Mb (202.74 Mb) of the cacao genome (Table 1; Figure 2). This coverage corresponds to 62.02% of the total available sequence (326.90 Mb), which covers 76.02% of the estimated genome of the B97-61/B2 genotype (430 Mb). Of the 10 chromosomes (Chrs) from the physical “*in silico*” map constructed in the present study, eight Chrs showed proportions above 50% (Chr 6, Chr 7, Chr 5, Chr 2, Chr 9, Chr 3, Chr 4 and Chr 1) and three Chrs showed proportions above 70% (Chr 3 and Chr 1) of the B97-61/B2 genotype sequenced chromosomes (Table 1).

**Figure 2.**
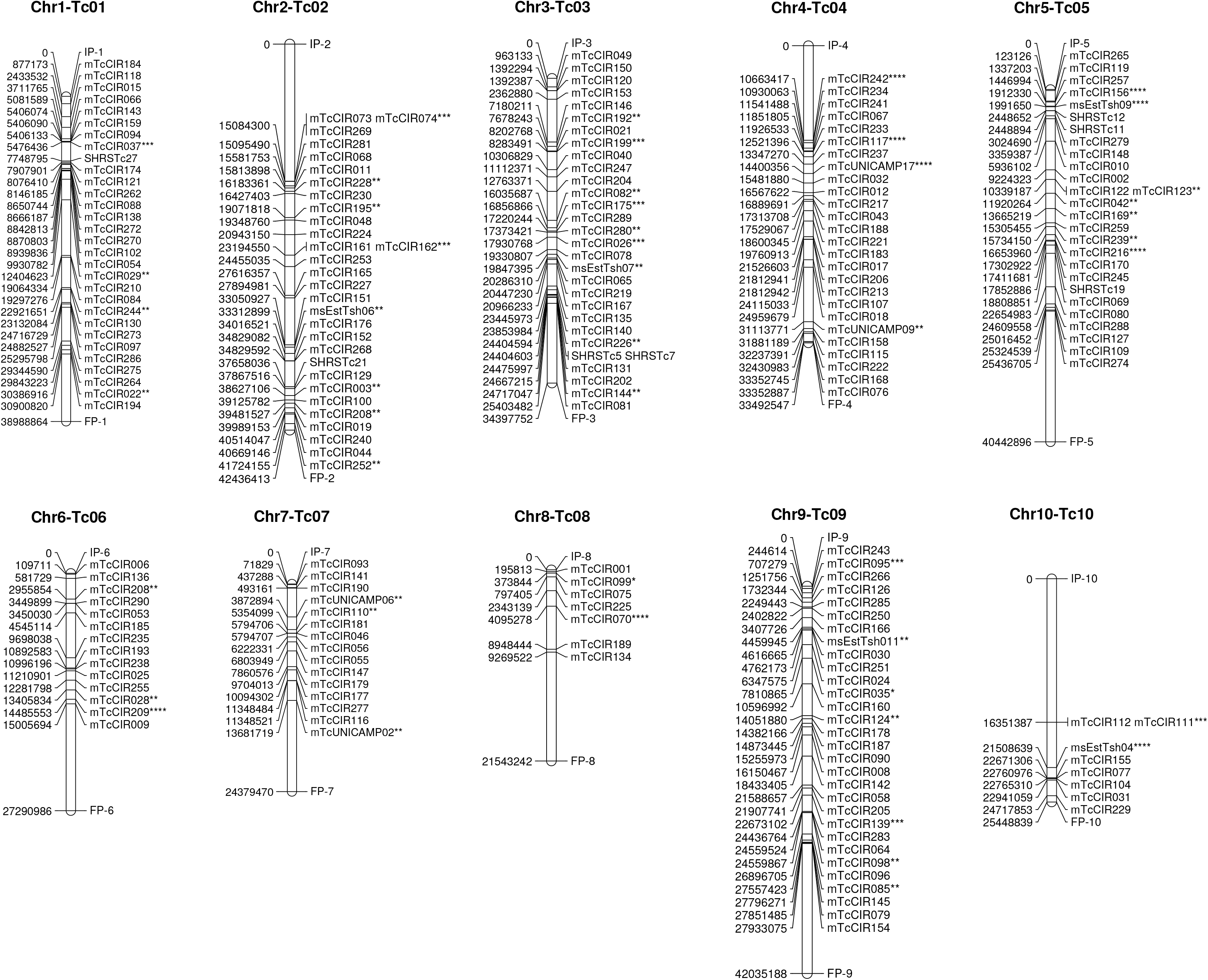
Physical “*in silico*” map constructed for the cacao tree (*Theobroma cacao* L.) using the sequence of the polymorphic markers detected for the 265 individuals of an F1 segregating population (TSH 1188 X CCN 51). This map consists of 217 SSRs covering more than 200 Mb (202.74 Mb) of the available cacao genome. The asterisks shown at the end of the name of some markers represent the following: *, deviation of segregation (ds); **, not aligned in the multipoint genetic linkage map; ***, not aligned in the same linkage group of the multipoint genetic map; ****, (ds) and not aligned in the multipoint genetic map; and *****, (ds) and not aligned in the same linkage group of the multipoint genetic map. IP and FP represent the initial and final positions of the chromosomes, respectively, obtained from the available cacao genome.

Of the 13 deviated markers from the F1 segregation patterns, 11 markers (84.62%) were allocated to the physical map. Moreover, this map positioned 38 SSRs that were not associated with any LGs of the multipoint genetic map (Figure 2). However, the multipoint genetic map of the present study accounted for 9 SSRs that were not aligned to any chromosomes of the physical map (Figure 1), demonstrating that both strategies were important for genome characterization. The distribution of the number of markers across the LGs and Chrs was similar between the linkage and physical maps (Table 1).

### Phenotypic Evaluation of Black Pod Disease

A total of 265 individuals from the F1 segregating population, parental clones TSH 1188 and CCN 51, and cultivars SCA 6 (resistant) and Catongo (susceptible) were used to evaluate the responses to black pod infection caused by the three *Phytophthora* species. The parental clones and cultivars were used as references (checks) in the trials, allowing for the estimation of residual effects, and together with the F1 individuals, the investigation of genetic variances, covariances and correlations. The strategy used here was to test different structures of variances and covariances for the genetic-effects matrix grouping in the trials or boxes within each trial (Supplementary Table A).

For both *P. palmivora* and *P. capsici*, the models that better explained the genetic variation were compound symmetry heterogeneous (CS_Het_) for the trials and factor analytic of order 1 (FA) for the boxes within each trial. For *P. citrophthora*, the diagonal (DIAG) model was better adjusted for both trials and boxes within the trial. These results show that genetic variances between the trials and among the boxes within each trial were heterogeneous for the three *Phytophthora* species. However, genetic covariances and correlations were only detected for *P. palmivora* and *P. capsici* species, with an equal estimated correlation between the trials and different correlations among the boxes within each trial, varying according to the pairwise combination involving the boxes (Supplementary Table 2).

The residual effects were also tested for complex models (results not shown). Convergence was not reached for these complex models based on the three *Phytophthora* species, which indicated that most of the variability likely reflected random genetic effects or that the adjustment of all the variance and covariance structures was too complex to reach convergence. The Wald test for fixed effects showed that only the boxes were significant (p-value < 0.001) for the three *Phytophthora* species (results not shown).

Comparative analyses between the black pod reactions of the inoculated parental clones (TSH 1188 and CCN 51) and cultivars (SCA 6 and Catongo) used as references revealed the efficiency of these cultivars as resistance and susceptibility standards, respectively (Supplementary Table 3). The parental clone CCN 51 showed high resistance to *P. palmivora* and moderate resistance to both *P. citrophthora* and *P. capsici*, whereas TSH 1188 showed susceptibility to all species, when compared to the SCA 6 clone. The predicted mean of the F1 population was consistently higher than that of CCN 51 and SCA 6 and lower than that of TSH 1188 and Catongo. Phenotypic 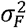 and genotypic 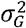 variances were observed in the F1 population for all species. Higher genotypic variance was observed for *P. citrophthora* (1.164), followed by *P. palmivora* (1.151) and *P. capsici* (0.848). High heritability (*h*^2^) was observed from the leaf-disc trials at 0.815 in *P. palmivora*, 0.903 in *P. citrophthora* and 0.639 in *P. capsici*. The coefficient of variation (CV %) varied from 6.757 to 8.511 (Supplementary Table A).

High and statistically significant Pearson correlation estimates were observed between *P. palmivora* and *P. citrophthora* (*r* = 0.730; p-value = 0.000) and between *P. citrophthora* and *P. capsici* (*r* = 0.630; p-value = 0.000), whereas a low and statistically significant Pearson correlation estimate was observed between *P. palmivora* and *P. capsici* (*r* = 0.340; p-value = 1.154 × 10^-8^).

### QTL Mapping

QTL mapping was performed based on the multipoint genetic map and predicted means of black pod disease from infections with *P. palmivora* (BP-Pp), *P. citrophthora* (BP-Pct) and *P. capsici* (BP-Pc). To identify QTLs, LOD score-based thresholds were obtained for the three *Phytophthora* species using 1,000 permutation tests, based on the method of Chen and Storey (2006), and produced the following values: 3.112 for BP-Pp, 3.058 for BP-Pct, and 3.174 for BP-Pc (Figure 3). A total of 6 QTLs were detected: 1 QTL for BP-Pp (LG 6), 1 QTL for BP-Pct (LG 6) and 4 QTLs for BP-Pc (LG 1, LG 2, LG 3 and LG 4) (Figure 3, Table 2, Supplementary Fig.2). Common QTLs were not detected among the three *Phytophthora* species, although the Pearson correlation estimates were high and statistically significant between BP-Pp and BP-Pct and between BP-Pct and BP-Pc.

**Figure 3.**
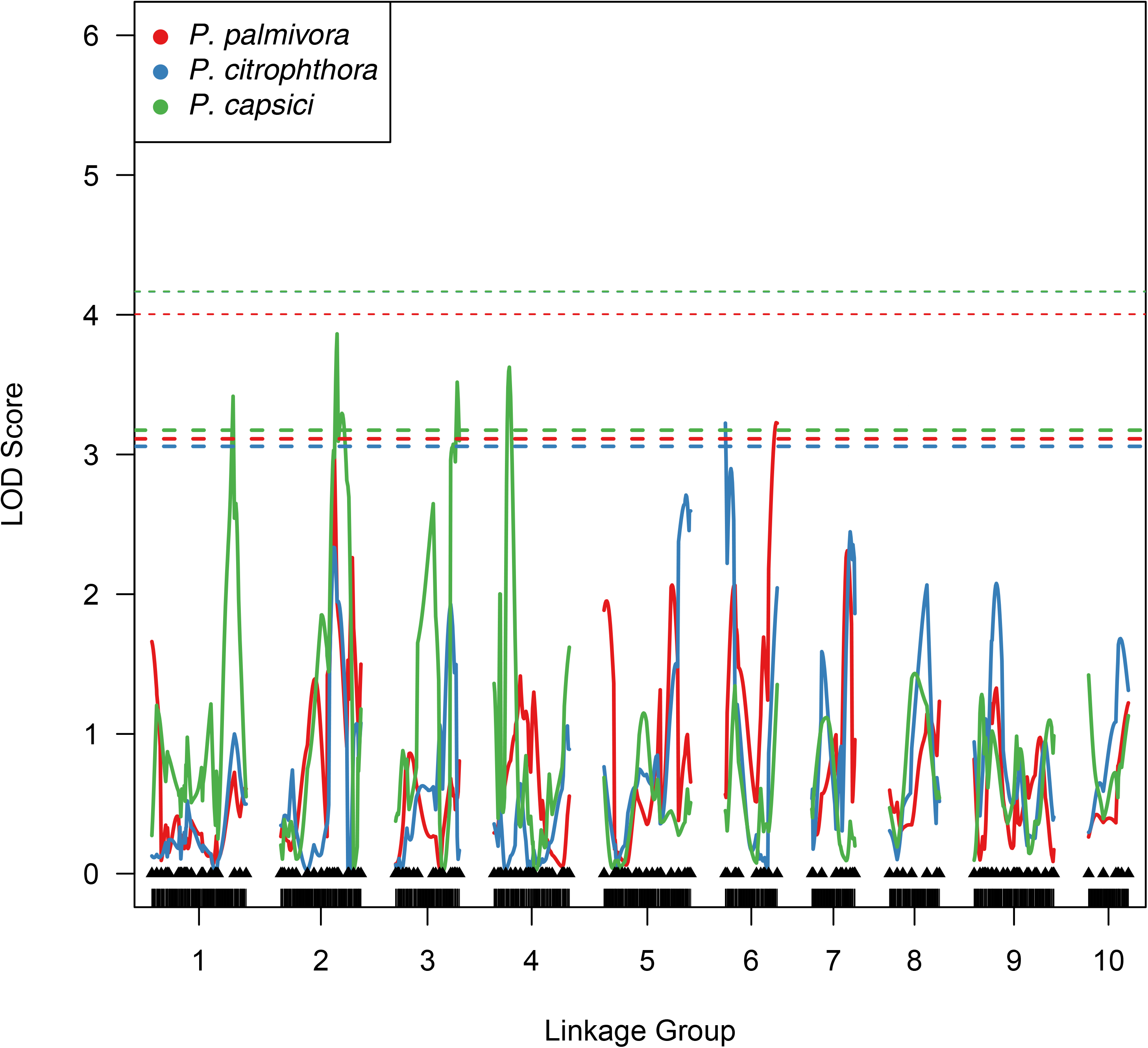
QTL mapping for the cacao tree (*Theobroma cacao* L.) associated with resistance to black pod disease resulting from infections with *P. palmivora*, *P. citrophthora* and *P. capsici*. The LOD score-based thresholds obtained from 1,000 permutation tests are plotted and are based on the method of Chen and Storey (2006) (thick dotted lines).

The proportion of phenotypic variation (R^2^) explained by the QTLs ranged from 1.77% to 3.29% (Table 2). The segregation patterns of the QTLs were 1:2:1 or 1:1. Of the 6 mapped QTLs, 5 QTLs (83.33%) had a significant additive effect for the parental clone TSH 1188 (i.e., with a LOD score higher than 0.834, that is the threshold with 1 degree of freedom and 5% error probability), 1 QTL (16.66%) had a significant additive effect for the parental clone CCN 51, and 2 QTLs (33.33%) had a significant dominance effect from the interaction between both parents. Three QTLs (50.00%) had only an additive effect for the parental clone TSH 1188, whereas the other three QTLs showed at least two different effects that explained the phenotypic variation. QTLs with a significant additive effect were not exclusively observed for the parental clone CCN 51 (Table 2).

**Table 2.**
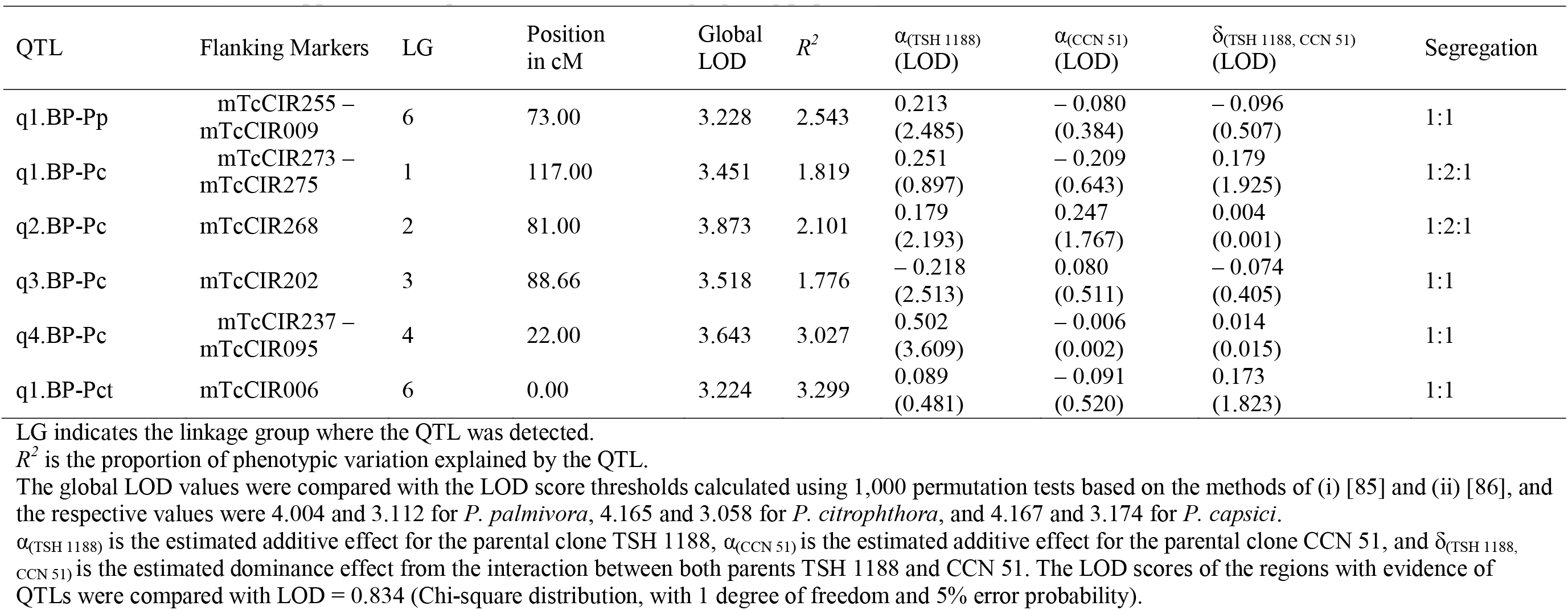
Effects of QTLs mapped to black pod disease for an F1 segregating population.

| QTL | Flanking Markers | LG | Position<br>in cM | Global<br>LOD | $R^2$ | $\alpha_{(TSH\ 1188)}$<br>(LOD) | $\alpha_{(CCN\ 51)}$<br>(LOD) | $\delta_{(TSH\ 1188,\ CCN\ 51)}$<br>(LOD) | Segregation |
| --- | --- | --- | --- | --- | --- | --- | --- | --- | --- |
| q1.BP-Pp | mTcCIR255 –<br>mTcCIR009 | 6 | 73.00 | 3.228 | 2.543 | 0.213<br>(2.485) | – 0.080<br>(0.384) | – 0.096<br>(0.507) | 1:1 |
| q1.BP-Pc | mTcCIR273 –<br>mTcCIR275 | 1 | 117.00 | 3.451 | 1.819 | 0.251<br>(0.897) | – 0.209<br>(0.643) | 0.179<br>(1.925) | 1:2:1 |
| q2.BP-Pc | mTcCIR268 | 2 | 81.00 | 3.873 | 2.101 | 0.179<br>(2.193) | 0.247<br>(1.767) | 0.004<br>(0.001) | 1:2:1 |
| q3.BP-Pc | mTcCIR202 | 3 | 88.66 | 3.518 | 1.776 | – 0.218<br>(2.513) | 0.080<br>(0.511) | – 0.074<br>(0.405) | 1:1 |
| q4.BP-Pc | mTcCIR237 –<br>mTcCIR095 | 4 | 22.00 | 3.643 | 3.027 | 0.502<br>(3.609) | – 0.006<br>(0.002) | 0.014<br>(0.015) | 1:1 |
| q1.BP-Pct | mTcCIR006 | 6 | 0.00 | 3.224 | 3.299 | 0.089<br>(0.481) | – 0.091<br>(0.520) | 0.173<br>(1.823) | 1:1 |

LG indicates the linkage group where the QTL was detected.
$R^2$ is the proportion of phenotypic variation explained by the QTL.
The global LOD values were compared with the LOD score thresholds calculated using 1,000 permutation tests based on the methods of (i) [85] and (ii) [86], and the respective values were 4.004 and 3.112 for *P. palmivora*, 4.165 and 3.058 for *P. citrophthora*, and 4.167 and 3.174 for *P. capsici*.
$\alpha_{(TSH\ 1188)}$ is the estimated additive effect for the parental clone TSH 1188, $\alpha_{(CCN\ 51)}$ is the estimated additive effect for the parental clone CCN 51, and $\delta_{(TSH\ 1188,\ CCN\ 51)}$ is the estimated dominance effect from the interaction between both parents TSH 1188 and CCN 51. The LOD scores of the regions with evidence of QTLs were compared with LOD = 0.834 (Chi-square distribution, with 1 degree of freedom and 5% error probability).

The QTL identified for *P. palmivora* (q1.BP-Pp) was located on LG 6 (73.00 cM) and explained 2.543% of the phenotypic variation (Table 2). This QTL showed a significant additive effect for the parental clone TSH 1188 and had a segregation ratio of 1:1. The QTL identified for *P. citrophthora* (q1.BP-Pct) was also located on LG 6 (0.00 cM) at position mTcCIR006 and explained 3.299% of the phenotypic variation. This QTL showed a significant dominance effect from the interaction between both parents and had a segregation ratio of 1:1. The 4 QTLs identified for *P. capsici* explained 9.889% of the phenotypic variation. The first QTL (q1.BP-Pc) was located on LG 1 (117.00 cM) and showed either an additive effect for TSH 1188 or a dominance effect from the interaction between TSH 1188 and CCN 51. The second QTL (q2.BP-Pc) was located on LG 2 (81.00 cM), at position mTcCIR268, and showed two significant additive effects—one for each parent. The third QTL (q3.BP-Pc) was located on LG 3 (88.66 cM), at position mTcCIR202, and had a significant additive effect for TSH 1188. The fourth (and last) QTL (q4.BP-Pc) was located on LG 4 (22.00 cM) and had an additive effect for TSH 1188. These 4 QTLs for *P. capsici* explained 1.819%, 2.101%, 1.776% and 3.027% of the phenotypic variation and exhibited segregation ratios of 1:2:1 (q1.BP-Pc and q2.BP-Pc) or 1:1 (q3.BP-Pc and q4.BP-Pc).

The highest LOD score peaks of the mapped QTLs were observed for BP-Pc (Table 2). We identified peak LOD scores of 3.873, 3.643, 3.518 and 3.451 for q2.BP-Pc, q4.BP-Pc, q3.BP-Pc and q1.BP-Pc, respectively. However, although the LOD scores of the mapped QTLs for BP-Pp and BP-Pct were lower (3.228 for q1.BP-Pp and 3.224 for q1.BP-Pct), the proportions of phenotypic variation were higher, with the exception of the QTL q1.BP-Pp (R^2^ = 2.543), when compared with the QTL q4.BP-Pc (R^2^ = 3.027).

### Co-localization of Disease Resistance-Related Genes and QTL Regions

We further investigated the genes located close to the QTL regions associated with the three *Phytophthora* species using the chromosomal locations of the markers as a reference. These chromosomal locations were obtained from the first assembled *T. cacao* L. genome (Argout et al., 2011), which is available in the *CocoaGen DB*. Based on an average interval of 100 Kb for the six regions containing the mapped QTLs, we observed that most of the QTLs were located in genomic regions containing resistance-related genes (Appendix C).

## Discussion

### Genetic Linkage Map

The level of polymorphisms detected in the present study was expected based on previous cacao mapping studies (Flament *et al*., 2001; Risterucci *et al*., 2003; Allegre *et al*., 2012). The expected segregation ratios of most of the polymorphic loci indicated a favorable scenario for the construction of an integrated genetic map based on the multipoint approach. A total of 13 (5.68%) SSRs deviated from the expected segregation ratios based on the Chi-square tests and Bonferroni correction. However, these markers were not discarded for the construction of the genetic map because these distortions were not strong enough to cause potential problems. In the final genetic linkage map, three of these markers (mTcCIR035, mTcCIR099, and mTcCIR191) were positioned in three different LGs and helped to obtain a more precise representation of the cacao genome. In previously published genetic maps, approximately 11% of the markers exhibited a distortion of segregation (twofold of the proportion detected here) (Flament *et al*., 2001; Brown *et al*., 2007; Allegre *et al*., 2012). The origin of these distortions remains unknown but has been suggested to result from sub-lethal gametophytic selection, gamete-sporophytic incompatibility in the cacao gene systems or even structural changes, although the latter has never been reported in cacao studies (Fouet *et al*., 2011).

The cacao genetic linkage map constructed in the present study showed 10 LGs. The cacao tree is a diploid species with 20 chromosomes (Davie, 1935); therefore, this map represented the haploid number of cacao chromosomes (Figure 1). The marker distribution among the LGs was not uniform, and several gaps (approximately 10 to 20 cM) were clearly observed in the multipoint genetic map, primarily in LGs 6, 7, 8 and 10, which has also been described in previous genetic maps of cacao (Brown *et al*., 2007; Allegre *et al*., 2012). Of these LGs, saturation was lower for LG 10, as has also been described in other mapping studies (Brown *et al*., 2007; Allegre *et al*., 2012). One possible explanation for the gaps is that either the recombination events or the mapped loci were not evenly distributed for certain LGs (Souza *et al*., 2013). These low-density markers might correspond to either highly homozygous regions that have a lower recombination frequency or centromeric regions.

The variation observed among cacao genetic linkage maps could partially reflect the use of different parental clones, progenies (F1, F2 or backcross), types and amounts of molecular markers, and algorithms to order and position the markers. We propose that the latter is crucial for obtaining more precise genetic linkage maps that may provide increased correspondence with the cacao genome. Our multipoint genetic map constructed from a cross between TSH 1188 and CCN 51 clones will be important for further studies on cacao breeding. These clones have contrasting agronomic traits associated with productivity and resistance to disease that are important traits for cacao breeding (with the SCA 6 clone serving as a standard resistance source).

### Physical “In Silico” Map

The physical “in silico” map constructed here provides a rational guideline for cacao-breeding programs for the characterization of selected clones and germplasm collections using mapped SSRs, thus providing information for recurrent genome alignment studies, which can be difficult to obtain in certain scenarios. The high correspondence between linkage and physical maps (Supplementary Fig.3) clearly shows the power of the multipoint approach to construct genetic maps. In the present study, the physical map accounted for SSRs that were not present in any LGs of the genetic map, thus generating important additional information about the genome. However, the multipoint genetic map also accounted for SSRs that did not align to any chromosomes of the physical map, showing that both strategies are powerful and should be used whenever possible in genetic studies of cacao.

### Phenotypic Evaluation of Black Pod Disease

Phenotypic analyses of black pod disease were performed using a mixed-model approach that allowed for the consideration of different scenarios for random genetic effects from the adjusted model, resulting in a model that can better explain experimental conditions and provide accurate predictions of the F1 individuals for QTL mapping. The findings of the present study suggest that more complex models provide more powerful explanations of the genetic variability of the *Phytophthora* species. The differentiated reaction of the cacao genotypes in response to *Phytophthora* spp. infection detected in this study was also observed in previous studies (Campêlo *et al*., 1982). Campêlo *et al*. (1982) investigated the response of ‘comum’ variety cacao pods to *Phytophthora* species infection and reported that isolates of *P. citrophthora* showed higher virulence, whereas *P. palmivora* and *P. capsici* isolates showed intermediate and lower virulence, respectively. The mean values of the leaf lesions caused by *P. citrophthora* are approximately two times higher than the mean values of the leaf lesions caused by *P. palmivora* in clones TSH 1188 and CCN 51 (Bahia *et al*., 2015). The moderate resistance of clone CCN 51 to *Phytophthora* spp. reflects the presence of genes potentially transferred from ICS 95 (Santos *et al*., 2007), a moderately resistant clone that is an ancestor present in the CCN 51 genealogy. However, the low resistance to *Phytophthora* spp. presented by the clone TSH 1188 in this study was probably because the black pod isolates were more virulent, i.e. isolates 692 (*P. capsici*), 1043 (*P. citrophthora*), and 2196 (*P. palmivora*), obtained from CEPEC *Phytophthora* collection. Our findings differ from the results showed by Bahia *et al*. (2015), who used the black pod isolates 62 (*P. citrophthora*) and 252 (*P. palmivora*).

The predicted mean of the F1 population was closer to the average of more resistant parental clones (i.e., CCN 51 clone) for both *P. citrophthora* and *P. capsici*. The high estimates of heritability for the three *Phytophthora* species demonstrated that black pod disease in cacao may be controlled by a few genes and may be minimally influenced by the environment, suggesting a high magnitude for the correlation between phenotypic and genotypic values. Furthermore, the coefficients of variation for the three *Phytophthora* species were suitable, considering that the trials were performed under controlled conditions in a small laboratory area. Thus, the black pod trials were adequately conducted and were experimentally appropriate.

### QTL Mapping

The present study is the first to use multipoint genetic linkage mapping (Wu et al., 2002a, b; Margarido et al., 2007; Tong et al., 2010), a mixed-model approach (Robinson, 2012) and CIM for an outbred population (Gazaffi et al., 2014) for cacao genetic mapping. The results detected here could provide important insights that will increase our current understanding of the cacao genome and the genetic architecture of black pod disease. The six QTLs detected in this study were based on the LOD score thresholds obtained by permutation testing based on the method of Chen and Storey (2006) in that the present study considers different LOD score peaks to construct the statistical distribution and declare the level of significance; we believe that the use of this more relaxed significance criterion showed more acceptable results for detecting QTLs associated with black pod disease.

All of the QTLs detected here were mapped to different regions of the cacao genome. Thus, common QTLs were not observed among the three *Phytophthora* species, and these findings implicate important research directions that should be pursued. First, QTLs were observed in 5 (LG 1, LG 2, LG 3, LG 4 and LG 6) out of the 10 LGs constituting the cacao genome, showing that 50% of the genome included resistance-related genes to black pod. Second, the absence of common QTLs detected for the *Phytophthora* species suggested that their mechanisms of resistance could also be specific. This result is interesting because different mechanisms that utilize different resistance proteins and metabolic pathways could be specifically investigated and described. Moreover, this information could be useful for specific marker-assisted selection programs and cacao genetic-breeding programs, depending on the interest and purpose of the study.

The proportion of phenotypic variation (R^2^) explained by the QTLs detected here ranged from 1.77% to 3.29% (Table 2). The QTL mapping of the present study was performed based on predicted means obtained from the mixed-model approach. Because the genetic effects were declared to be random in the mixed models, the predicted means were corrected by a *shrinkage* factor, which provides genetic effects that should be very close to the real values. Thus, we believe that the proportion of R^2^ explained by the QTLs in the present study was very reliable. Similar proportions of phenotypic variation have also been detected by other QTL studies that used mixed models for the phenotypic analyses in different plant species, such as the rubber tree (Souza *et al*., 2013) and the common bean (Oblessuc *et al*., 2014). However, a great proportion of phenotypic variation remains unexplained, assuming that the heritability of the black pod disease is high. One possible explanation is that the mapping population size used in the present study (265 F1 individuals) was not large enough to detect other possible QTLs with similar proportions of phenotypic variation compared with the mapped QTLs (Beavis, 1994; Bernardo, 2010). Another possible explanation is that interactions between the mapped QTLs and other possible QTLs (epistasis) may explain the high proportion of phenotypic variation. Epistasis effects should be important for black pod resistance because the molecular mechanisms of this disease seem to be very complex, as reported recently by Nyadanu et al. (2012).

### Co-localization of Disease Resistance-Related Genes and QTL Regions

The mapped QTLs detected in the present study indicate genomic regions that should be further exploited to generate more polymorphisms for future studies of QTL mapping or to investigate resistance-related candidate genes. Flanking markers of the mapped QTLs were located in proximity to important genomic regions (Appendix C). An important predicted ortholog of leucine rich repeat (LRR)-containing receptors was located in the q2.BP-Pc and q4.BP-Pc regions; these receptors detect specific pathogenic peptides that signal to Pelle-family kinases (Dievart and Clark, 2004) and play central roles in signaling during pathogen recognition for the subsequent activation of defense mechanisms and developmental control (Afzal et al., 2008). Two important genes encoding peroxidase 65 and superoxide dismutase were mapped in the q2.BP-Cp region; peroxidase genes are involved in the response to environmental stresses, such as wounds and pathogen attacks (Kawano, 2003), whereas superoxide dismutase acts as an essential component in defense mechanisms against oxidative stress and pathogens (Bowler et al., 1992). Genes assigned as Castor were identified in the q4.BP-Pc region; this gene is an ion channel that is likely involved in fungal entry into root epidermal cells during the establishment of mycorrhizal symbiosis (Charpentier et al., 2008). Genes potentially involved in pathogen defense were identified in the q1.BP-Pp region, such as CPR30 that acts as a negative regulator of plant defense responses (Gou et al., 2009), Cys5 that encodes a small protein with antimicrobial and antifungal activities that is expressed in various plant tissues (Lay and Anderson, 2005) and Y-3 that interacts with the kinase domain under various experimental conditions, suggesting that Y-3 may be involved in stress conditions, such as mechanical wounding and pathogen infection (Tarutani and Sasaki, 2004). Moreover, we identified genes associated with the stress response and programmed cell death, such as BAG5 [101] and UPL5 [102].

### Conclusion

The number of genes involved in resistance to *Phytophthora* diseases is comparable to that observed in other species. Furthermore, the common regions of certain QTLs across different genetic groups confirm the existence of and potential use of these genes for marker-assisted selection programs. The identification of multiple QTLs involved in resistance to *Phytophthora* may be particularly useful when transferring different QTLs into an elite clone using a marker-assisted selection scheme. However, it would be interesting to evaluate interaction of these genes with other agronomic traits of interest to determine the relationship between resistance phenotypes in the field and yield components. We propose that a detailed functional genomics study should be performed to confirm the roles of these QTLs associated with resistance to black pod disease. The multipoint genetic linkage and physical maps constructed in the present study will be useful for these further analyses.

## Acknowledgments

The authors would like to thank the Comissão Nacional de Desenvolvimento Científico e Tecnológico (CNPq), Fundação de Amparo à Pesquisa do Estado da Bahia (FAPESB), Fundação de Amparo à Pesquisa do Estado de São Paulo (FAPESP, 2008/53197-4), Banco do Nordeste Brasileiro (BNB), Comissão Executiva do Plano da Lavoura Cacaueira (CEPLAC), Universidade Estadual de Campinas (UNICAMP), Universidade Estadual de Santa Cruz (UESC), and Coordenação de Aperfeiçoamento de Pessoal de Nível Superior (PROCAD-UESC, Computational Biology Program) for the grants provided for this research, FAPESP for the PhD fellowship to CBCS (2010/0354-2, 2012/11109-0), and CNPq for the PhD fellowships to MAB, JRBFR and ESLS and for the research fellowships awarded to AAFG and APS. The authors would also like to thank the researchers from UNICAMP, UESC, MCCS and CEPLAC for supplying the physical structure required to perform this study. Additionally, the authors would like to thank Fernanda Bispo and Ademilde Cerqueira for assistance with black pod phenotyping, and Dr. Karina Gramacho for her invaluable help during the correction of the manuscript.

## References

Afzal AJ, Wood AJ, Lightfoot DA, 2008. Plant receptor-like serine threonine kinases: roles in signaling and plant defense. Molecular Plant-Microbe Interactions 21, 507–17.

Akaike H, 1974. A new look at the statistical model identification. IEEE Transactions on Automatic Control 19, 716–23.

Allegre M, Argout X, Boccara M et al., 2012. Discovery and mapping of a new expressed sequence tag-single nucleotide polymorphism and simple sequence repeat panel for large-scale genetic studies and breeding of *Theobroma cacao* L. DNA Research 19, 23–35.

Argout X, Salse J, Aury J-M et al., 2011. The genome of *Theobroma cacao*. Nature Genetics 43, 101–8.

Bahia RDC, Aguilar-Vildoso CI, Luz EDMN, Lopes UV, Machado RCR, Corrêa RX, 2015. Resistance to black pod disease in a segregating cacao tree population. Tropical Plant Pathology 40, 13–8.

Barreto MA, Santos JCS, Corrêa RX, Luz EDMN, Marelli J, Souza AP, 2015. Detection of genetic resistance to cocoa black pod disease caused by three Phytophthora species. Euphytica 206, 677–87.

Bowler C, Montagu MV, Inze D, 1992. Superoxide dismutase and stress tolerance. Annual Review of Plant Physiology and Plant Molecular Biology 43, 83–116.

Boza EJ, Motamayor JC, Amores FM et al., 2014. Genetic characterization of the cacao cultivar CCN 51: its impact and significance on global cacao improvement and production. Journal of the American Society for Horticultural Science 139, 219–29.

Brown JS, Phillips-Mora W, Power EJ et al., 2007. Mapping QTLs for resistance to frosty pod and black pod diseases and horticultural traits in *Theobroma cacao*. Crop Science 47, 1851–8.

Campêlo A, Luz E, Resnik F, 1982. Podridao parda do cacaueiro no Estado da Bahía, Brasil, 1: virulencia das espécies de Phytophthora. Revista Theobroma 12, 1–6.

Charpentier M, Bredemeier R, Wanner G, Takeda N, Schleiff E, Parniske M, 2008. *Lotus japonicus* CASTOR and POLLUX are ion channels essential for perinuclear calcium spiking in legume root endosymbiosis. Plant Cell 20, 3467–79.

Chen L, Storey JD, 2006. Relaxed significance criteria for linkage analysis. Genetics 173, 2371–81.

Clair DA, 2010. Quantitative disease resistance and quantitative resistance loci in breeding. Annual Review of Phytopathology 48, 247–68.

Clement D, Risterucci AM, Motamayor JC, N’goran J, Lanaud C, 2003. Mapping QTL for yield components, vigor, and resistance to *Phytophthora palmivora* in *Theobroma cacao* L. Genome 46, 204–12.

Crouzillat D, Ménard B, Mora A, Phillips W, Pétiard V, 2000. Quantitative trait analysis in *Theobroma cacao* using molecular markers. Euphytica 114, 13–23.

Davie JH, 1935. Chromosome studies in the malvaceae and certain related families. II. Genetica 17, 487–98.

Flament MH, Kebe I, Clement D et al., 2001. Genetic mapping of resistance factors to *Phytophthora palmivora* in cocoa. Genome 44, 79–85.

Gazaffi R, Margarido GRA, Pastina MM, Mollinari M, Garcia AAF, 2014. A model for quantitative trait loci mapping, linkage phase, and segregation pattern estimation for a full-sib progeny. Tree Genetics & Genomes 10, 791–801.

Gou M, Su N, Zheng J et al., 2009. An F-box gene, CPR30, functions as a negative regulator of the defense response in Arabidopsis. Plant Journal 60, 757–70.

Guest D, 2007. Black pod: diverse pathogens with a global impact on cocoa yield. Phytopathology 97, 1650–3.

Henderson CR, Kempthorne O, Searle SR, Von Krosigk C, 1959. The estimation of environmental and genetic trends from records subject to culling. Biometrics 15, 192–218.

ICCO, 2014. Annual report. London United Kingdon 2013-2014. The International Cocoa Organization. [http://www.icco.org/]. Accessed 5 April 2014.

CGD, 2015. International Cocoa Germoplasm Database - CGD. [http://www.icgd.rdg.ac.uk/ref_search.php?refcode=SHR96A]. Accessed 14 December 2016.

Kawano T, 2003. Roles of the reactive oxygen species-generating peroxidase reactions in plant defense and growth induction. Plant Cell Reports 21, 829–37.

Kosambi DD, 1943. The estimation of map distances from recombination values. Annals of Eugenics 12, 172–5.

Kover PX, Caicedo AL, 2001. The genetic architecture of disease resistance in plants and the maintenance of recombination by parasites. Molecular Ecology 10, 1–16.

Lanaud C, Fouet O, Clément D et al., 2009. A meta–QTL analysis of disease resistance traits of *Theobroma cacao* L. Molecular Breeding 24, 361–74.

Lanaud C, Risterucci AM, Pieretti I, N’goran JaK, Fargeas D, 2004. Characterisation and genetic mapping of resistance and defence gene analogs in cocoa (*Theobroma cacao* L.). Molecular Breeding 13, 211–27.

Lay FT, Anderson MA, 2005. Defensins-components of the innate immune system in plants. Current Protein and Peptide Science 6, 85–101.

Lopes UV, Monteiro WR, Pires JL, Clement D, Yamada MM, Gramacho KP, 2011. Cacao breeding in Bahia, Brazil -strategies and results. Crop Breeding and Applied Biotechnology 1, 73–81.

Luz EDMN, Sgrillo RB, Santosfilho LP, 2004. Estimativas de danos e perdas causadas por doenças no cacaueiro. In: Proceedings of the Proceedings of the WorkShop de Epidemiologia de doença. Viçosa, 67–79.

Magalhães DMA, Luz EDMN, Lopes UV, Niella ARR, Damaceno VO, 2016. Leaf disc method for screening Ceratocystis wilt resistance in cacao. Tropical Plant Pathology 41, 155–61.

Margarido GR, Souza AP, Garcia AA, 2007. OneMap: software for genetic mapping in outcrossing species. Hereditas 144, 78–9.

Michelmore RW, 2003. The impact zone: genomics and breeding for durable disease resistance. Current Opinion in Plant Biololy 6, 397–404.

Motamayor JC, Mockaitis K, Schmutz J et al., 2013. The genome sequence of the most widely cultivated cacao type and its use to identify candidate genes regulating pod color. Genome Biology 14, r53.

Nyadanu D, Akromah R, Adomako B et al., 2012. Biochemical mechanisms of resistance to black pod disease in cocoa (*Theobroma cacao* L.). American Journal of Biochemistry and Molecular Biology 3, 20–37.

Nyassé S, Cilas C, Herail C, Blaha G, 1995. Leaf inoculation as an early screening test for cocoa (*Theobroma cacao* L.) resistance to Phytophthora black pod disease. Crop Protection 14, 657–63.

Nyassé S, Mousseni IBE, Eskes AB, 2003. Selection for resistance to black pod disease and yield gains prediction by use of selected cocoa varieties in Cameroon. Plant Genetic Resources 1, 157–60.

Oblessuc PR, Baroni RM, Da Silva Pereira G et al., 2014. Quantitative analysis of race-specific resistance to *Colletotrichum lindemuthianum* in common bean. Molecular Breeding 34, 1313–29.

Oliveira ML, Luz EDMN, 2005. Identificação e Manejo das Principais Doenças do Cacaueiro no Brasil. Ilhéus: CEPLAC/CEPEC/SEFIT.

Patterson HD, Thompson R, 1971. Recovery of inter-block information when block sizes are unequal. Biometrika 58, 545–54.

Payne RW, Murray DA, Harding SA, Baird DB, Soutar DM, 2010. GenStat for Windows, 13th Edn. Introduction. Hemel Hempstead, UK: VSN International.

Rehem BC, Almeida AA, Correa RX, Gesteira AS, Yamada MM, Valle RR, 2010. Genetic mapping of *Theobroma cacao* (Malvaceae) seedlings of the Parinari series, carriers of the lethal gene Luteus-Pa. Genetics and Molecular Research 9, 1775–84.

Risterucci AM, Paulin D, Ducamp M, N’goran JA, Lanaud C, 2003. Identification of QTLs related to cocoa resistance to three species of Phytophthora. Theoretical and Applied Genetics 108, 168–74.

Robinson GK, 2012. That BLUP is a good thing: the estimation of random effects. Statistical Science 6, 15–32.

Santos RMF, Lopes UV, Bahia RDC, Machado RCR, Ahnert D, Corrêa RX, 2007. Marcadores microssatélites relacionados com a resistência à vassoura-de-bruxa do cacaueiro. Pesquisa Agropecuária Brasileira 42, 1137–42.

Santos ES et al., 2009. Identifi cação de resistência genética do cacaueiro à podridão-parda. Pesq. agropec. bras., 44(4), p.413–416.

Schultes RE, Cuatrecasas J, 1964. Cacao and its allies. A taxonomic revision of the genus *Theobroma*. Review. Economic Botany Journal 19, 416–7.

Souza LM, Gazaffi R, Mantello CC et al., 2013. QTL mapping of growth-related traits in a full-sib family of rubber tree (*Hevea brasiliensis*) evaluated in a sub-tropical climate. PLoS One 8, e61238.

Tahi GM, Kébé BI, Sangare A, Mondeil F, Cilas C, Eskes AB, 2006. Foliar resistance of cacao (*Theobroma cacao*) to *Phytophthora palmivora* as an indicator of pod resistance in the field: interaction of cacao genotype, leaf age and duration of incubation. Plant Pathology 55, 776–82.

Tarutani Y, Sasaki A, 2004. Identification of three clones which commonly interact with the kinase domains of highly homologous two receptor-like kinases, RLK902 and RKL1. Bioscience, Biotechnology, and Biochemistry 68, 2581–7.

Tong C, Zhang B, Shi J, 2010. A hidden Markov model approach to multilocus linkage analysis in a full-sib family. Tree Genetics & Genomes 6, 651–62.

Voorrips RE, 2002. MapChart: software for the graphical presentation of linkage maps and QTLs. Journal of Heredity 93, 77–8.

Wu R, Ma CX, Painter I, Zeng ZB, 2002a. Simultaneous maximum likelihood estimation of linkage and linkage phases in outcrossing species. Theoretical Population Biology 61, 349–63.

Wu R, Ma C-X, Wu SS, Zeng Z-B (2002b) Linkage mapping of sex-specific differences. Genet Res 79:85–96. doi: 10.1017/S0016672301005389.

